# Disinhibitory circuitry gates associative synaptic plasticity in olfactory cortex

**DOI:** 10.1101/2020.09.24.312686

**Authors:** Martha Canto-Bustos, F. Kathryn Friason, Constanza Bassi, Anne-Marie M. Oswald

## Abstract

Inhibitory microcircuits play an essential role in regulating cortical responses to sensory stimuli. Interneurons that inhibit dendritic or somatic integration in pyramidal neurons act as gatekeepers for neural activity, synaptic plasticity and the formation of sensory representations. Conversely, interneurons that specifically inhibit other interneurons can open gates through disinhibition. In the rodent piriform cortex, relief of dendritic inhibition permits long-term potentiation (LTP) of the recurrent synapses between pyramidal neurons (PNs) thought to underlie ensemble odor representations. We used an optogenetic approach to identify the inhibitory interneurons and disinhibitory circuits that regulate LTP. We focused on three prominent inhibitory neuron classes-somatostatin (SST), parvalbumin (PV), and vasoactive intestinal polypeptide (VIP) interneurons. We find that LTP is gated by the inactivation SST or PV interneurons or by activation of VIP interneurons. Further, activation of VIP interneurons strongly inhibits putative SST-cells during LTP induction, but only weakly inhibit PV-interneurons. Taken together, these findings suggest that VIP-interneurons mediate a disinhibitory circuit that can regulate synaptic plasticity during olfactory processing.

---

Throughout the cortex, the response properties of individual neurons as well as coordinated ensemble activity are refined by sensory experience. One underlying feature of experience-dependent plasticity is long-term changes in synaptic strength within cortical circuits. While the mechanisms underlying excitatory synaptic plasticity have been extensively studied (Abbott and Nelson 2000, Malenka and Bear 2004), less is known about the role cortical circuitry plays in gating changes in synaptic strength. Inhibitory interneurons regulate both dendritic integration and neural activity, two major factors in synaptic plasticity. Thus, inhibitory circuits can play a key role in the enhancement of synaptic connections (Artinian and Lacaille 2018, Lucas and Clem 2018). In this study, we elucidate the inhibitory and disinhibitory circuit motifs that gate synaptic plasticity at recurrent excitatory synapses in the olfactory cortex.

The anterior piriform cortex (APC) processes olfactory information, and performs both sensory and associative cortical functions. Located two synapses from the periphery; the APC is a primary cortical area representing odor inputs. Olfactory receptor neurons in the nose project to Mitral and Tufted (M/T) neurons in the olfactory bulb (OB)(Mombaerts, Wang et al. 1996). M/T neurons then project directly to APC and synapse with pyramidal neurons (PNs)(Haberly and Price 1977). Unlike other primary sensory cortices, APC lacks a topological representation of odor identity. M/T axons project diffusely and randomly (Sosulski, Bloom et al. 2011, Igarashi, Ieki et al. 2012) to activate distributed neural ensembles (Illig and Haberly 2003, Rennaker, Chen et al. 2007, Stettler and Axel 2009). Distributed odor representations are further supported by uniform intracortical excitatory connectivity across the APC (Franks, Russo et al. 2011). It is postulated that odor-specific ensembles are constructed by strengthening excitatory synapses between pyramidal neurons co-activated by odor components (Haberly 2001, Wilson and Sullivan 2011). In support of this hypothesis, intracortical synapses between PNs are strengthened following odor learning *in vivo* (Saar, Grossman et al. 2002) and through pairing of afferent and intracortical stimulation *in vitro* (Kanter and Haberly 1993, Johenning, Beed et al. 2009). Hence, APC circuitry also supports early stage, associative odor processing.

The APC is an ideal structure for investigating the circuit and synaptic plasticity mechanisms that underlie sensory representations. PNs receive compartmentalized excitatory inputs on their apical dendrites from two distinct fiber tracts. M/T cell afferents synapse distally, while PN axons form a proximal intracortical fiber tract (Haberly and Price 1977, Haberly and Price 1978). Co-activation of afferent and intracortical fiber tracts strengthens intracortical synapses onto PNs through NMDA receptor (R) dependent, associative LTP (Kanter and Haberly 1993). Further, NMDAR EPSPs and associative LTP induction are facilitated by dendritic disinhibition by GABA_A_ receptor antagonists (Kanter and Haberly 1993, Kanter, Kapur et al. 1996). While these studies suggest interplay between dendritic inhibition and disinhibition gates LTP induction at intracortical synapses, the inhibitory circuitry involved has not been identified.

Olfactory stimuli recruit both feedforward and recurrent inhibition onto PNs in APC (Poo and Isaacson 2009, Poo and Isaacson 2011). Feedforward inhibition is comparatively weak and diminishes with high frequency stimulation of the afferent pathway, whereas recurrent inhibition is strong, and increases with stimulation through synaptic facilitation and PN recruitment (Stokes and Isaacson 2010, Suzuki and Bekkers 2010a, Large, Vogler et al. 2016). Recurrent inhibition is mediated by inhibitory interneurons that express somatostatin (SST-INs) or parvalbumin (PV-INs) (Stokes and Isaacson 2010, Suzuki and Bekkers 2010a, Suzuki and Bekkers 2010b, Large, Kunz et al. 2016). SST-INs inhibit PN apical dendrites proximal to the soma, and are optimally located to regulate plasticity of intracortical synapses (Suzuki and Bekkers 2010b, Large, Kunz et al. 2016). PV-INs regulate spike activity and could also impact LTP through backpropagation (Johenning, Beed et al. 2009). A third class of interneurons-vasoactive intestinal polypeptide interneurons (VIP-INs); inhibit SST and PV-INs and could disinhibit pyramidal neurons (Lee, Kruglikov et al. 2013, Pfeffer, Xue et al. 2013, Karnani, Jackson et al. 2016). VIP-INs are a prominent in APC (Suzuki and Bekkers 2010b) but their inhibitory connections and function are unknown. We investigated the connectivity and functional roles of SST, PV and VIP-INs in APC. We find that activation of VIP-INs as well as inactivation of SST-INs or PV-INs promote LTP of intracortical synapses. VIP-INs strongly inhibit SST-INs during LTP induction, but only weakly inhibit PV-INs and PNs. Our finding suggests a VIP->SST>-PN disinhibitory circuit gates associative LTP in APC.

## Methods

### Mice

VIP-Cre (B6:Viptm1(cre)Zjh/J), SST-Cre (B6:Sst<tm2.1(cre)Zjh>/J) and PV-Cre mice express cre-recombinase (Taniguchi, He et al. 2011). These mice were crossed with Ai32 mice (B6;129S-Gt ROSA)26Sortm32 (CAG-COP4*H134R/EYFP)Hze/J) to express channelrhodopsin (ChR2) or Ai35 mice (B6.129S-*Gt(ROSA*)26Sortm35.1(*CAG-aop3/GFP)Hze*/J) to express archaerhodopsin (Arch) (Madisen, Mao et al. 2012). All mice are from Jackson Laboratory. All animals were bred, handled and treated in manner that was evaluated and approved by the Animal Care and Use Committee at the University of Pittsburgh IACUC, protocol #17070877.

### Slice preparation

APC brain slices were prepared from mice aged P19-35. The mice were anesthetized with isoflurane and the brain was removed and immersed in ice cold oxygenated (95% O_2_-5% CO_2_) ACSF (in mM: 125 NaCl, 2.5 KCl, 25 NaHCO_3_, 1.25 NaH_2_PO_4_, 1.0 MgCl_2_, 25 Glucose, 2.5 CaCl_2_) (all chemicals from Sigma, USA unless otherwise stated). Parasagittal slices (300 μm) were cut on a vibratome (Leica Biosystems) in ice-cold ACSF. The slices were transferred to warm ACSF (36°C) for 30 min, then 20-22°C for 1 hour, and recorded at 25-28°C.

### Electrophysiology

Recordings were performed using a MultiClamp 700B amplifier (Molecular Devices, Union City, CA). Data were low pass filtered (4 kHz) and digitized at 10 kHz using an ITC-18 (Instrutech) controlled by custom software (Recording Artist, https://bitbucket.org/rgerkin/recording-artist) written in IgorPro (Wavemetrics). Recording pipettes (4-10 MΩ) were pulled from borosilicate glass (1.5 mm, outer diameter) on a Flaming/Brown micropipette puller (Sutter Instruments). The series resistance (<20 MΩ) was not corrected. For PSPs the intracellular solution consisted of (in mM) 130 K-gluconate, 5 KCl, 2 MgCl_2_, 4 ATP-Mg, 0.3 GTP, 10 HEPES, and 10 phosphocreatine, 0.05% biocytin. For IPSC recordings, Qx-314 was added to the K-gluconate internal (holding potential 0 mV) or Cs-Glu-Qx solution was used (in mM, 130 Cs-Gluconate, 5 KCl, 2 MgCl_2_, 4 Mg-ATP, 0.3 GTP, 10 HEPES, 10 Phosphocreatine, 1 Qx-314, holding potential +30 mV). Neurons were visualized using infrared-differential interference contrast microscopy (IR-DIC, Olympus). For all recorded neurons, subthreshold response properties were obtained using a series of hyperpolarizing and depolarizing current steps (−50 pA to 50 pA, 1 s duration). We used specific criteria for identification of putative PV and SST interneurons based on intrinsic properties of identified PV and SST neurons in APC (see (Large, Kunz et al. 2016, for details). Neural identity was confirmed post hoc using intrinsic properties and biocytin fills.

### LTP induction

Electrical stimulation was delivered using concentric bipolar electrodes (FHC). The electrodes were placed in the LOT (L1a) and the L1b/L2 border. Stimuli (100 µs pulse width) were delivered through a stimulus isolation unit. Theta burst stimulation (TBS) of L1a consisted of 10 bursts of 4 pulses (100 Hz) delivered at 250 ms intervals (Kanter and Haberly 1993). TBS stimulation intensity was set near spike threshold for the recorded neuron. L1b was stimulated with a single weak pulse delivered between the 3^rd^ and 4^th^ pulse of each burst. Stimulation intensity was <30% of the maximum subthreshold EPSP (∼1-6 mV). Pre and post induction test pulses were delivered to L1b every 30 s. Baseline was collected for ∼5 min and LTP was induced within 10 min of patching the neuron. Input resistance was monitored throughout and neurons with deviations greater than 20% from baseline were excluded from analysis (n=10 neurons). We also monitored membrane potential throughout recording. Small fluctuations <5 mV were offset with constant current to maintain driving force. However, neurons with drift greater than 5 mV or exhibiting unstable membrane potential fluctuations were excluded from analysis (n=8).

### Pharmacology

The GABA_A_ receptor antagonist, gabazine (GZ, 20 µM in ACSF) was loaded into a regular patch pipette and focally applied to L1b using a gentle positive pressure (<1-5 s duration) see. Slices were oriented such that bath flowed from the soma to dendrite of the pyramidal cell to maintain GZ in the region of the dendrite. Every effort was made to ensure stability of GZ application including supplemental applications. In a subset of slices (n=5) from mice that expressed ChR2 in SST-INs, GZ application to L1b diminished optically evoked SST-mediated inhibition onto pyramidal cells (n=5). We verified that optically evoked IPSPs (Control: 6.3 ± 0.69 mV) in these slices were stably diminished by Gabazine application throughout the LTP protocol and did not differ between baseline (IPSP GZ: 0.89 ± 0.27 mV) and post-induction test pulses (IPSP GZ: 0.92 ± 0.28, p>0.05 WSR). In additional experiments, the NMDA receptor antagonist, DL-2-Amino-5-phosphonopentanoic acid (DL-APV, 10 mM, Sigma-Aldrich) was bath-applied as indicated in the main text.

### Optogenetic Stimulation

Shutter controlled full field stimulation with blue (473 nm) or green (520 nm) light (Prior) was delivered through the epifluorescence pathway of the microscope (Olympus) using a water-immersion objective (40x). Light intensity (5-10 mW) was adjusted to induce spike responses (ChR2 activation) or spike suppression (Arch inactivation). Light pulse duration varied by experiment as indicated in the main text.

### EPSP Analysis

EPSPs were analyzed using custom software written in IgorPro (Wavemetrics). Because we do not block inhibition during L1b test-pulses, we set narrow criteria to ensure that only the excitatory portion of mixed EPSP-IPSPs was analyzed. Recurrent inhibition is disynaptic and IPSPs are delayed with respect to excitation. Only EPSP onsets within 5 ms of the stimulus artifact were analyzed. Across all conditions the average EPSP onset was 3.7 ± 0.41 ms. EPSP peak amplitude was taken as the max amplitude within 5 ms of EPSP onset. We also measured the slope of the rising phase of the EPSP from onset to 80% of peak amplitude to further isolate the EPSP. Our core findings did not differ between peak and slope measurements.

### Statistics

All data is presented as mean ± SE. Statistical tests were performed using two tailed, one or two-sample, paired or unpaired Student’s t-test as appropriate. In cases of small sample sizes non-parametric tests were used, including the Mann-Whitney U-test (MWU) for unpaired data and the Wilcoxon Signed Ranks test (WSR) for paired data. For multiple comparisons we used ANOVA with post hoc Tukey Test (ANOVA).

## Results

We investigated the roles of SST, PV and VIP-INs, in gating associative LTP at intracortical excitatory synapses onto PNs. Afferent input to PNs arrives via L1a on distal L2 PN dendrites, while the intracortical fiber tract (L1b) is proximal (**Fig 1A**). L1a and L1b were easily identified under IR-DIC and independently stimulated using bipolar electrodes. Associative LTP is induced by pairing L1a and L1b stimulation using a theta burst stimulation (TBS) protocol (Kanter and Haberly 1993) consistent with respiration coupled M/T spike frequencies (Kepecs, Uchida et al. 2007, Carey and Wachowiak 2011). Briefly, strong TBS of L1a was paired with weak, single pulse stimulation of L1b (see methods, **Fig 1A**). This L1a+L1b pairing is hereafter denoted *induction*. L1a-TBS evoked low PN firing rates ranging from 0-8 Hz and L1b EPSPs ranged from 1-6 mV. Pre and post induction, L1b stimulation was delivered every 30 s. To avoid drift in recording integrity, potentiation was quantified as the average L1b EPSP amplitude 25-30 min following induction versus average baseline EPSP amplitude (5 min prior to induction).

**Figure 1:**
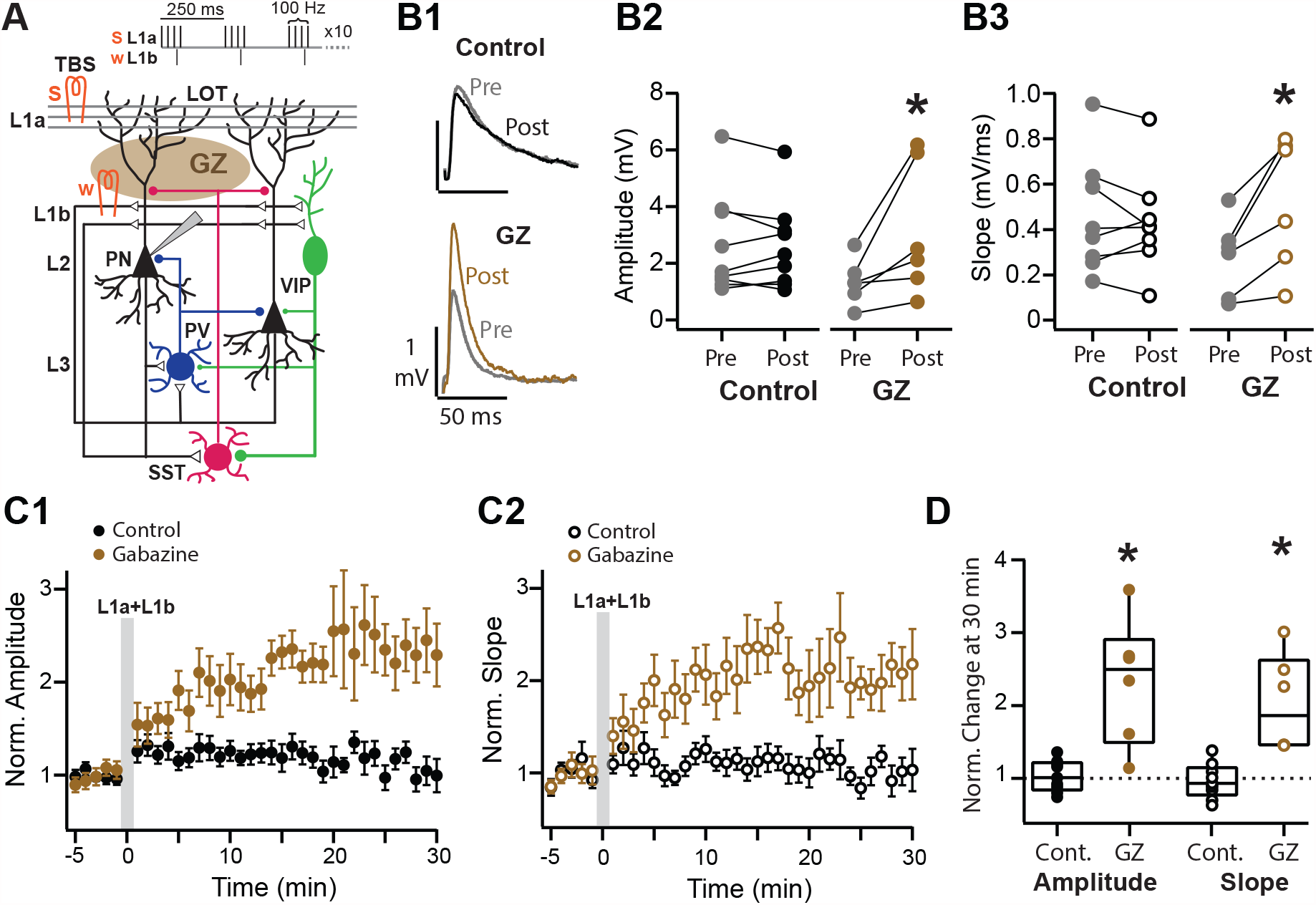
Associative LTP of intracortical synapses is gated by dendritic disinhibition. **A)** Schematic of APC circuit and stimulation paradigm. Strong (s) TBS of afferents (L1a) is paired with weak (w) single-pulses at the intracortical pathway (L1b). Gabazine (GZ) is focally applied to the dendrite region in L1b (tan oval). B1) Representative average traces pre (gray) and post induction in control (black) and with GZ(Tan). B2) Average EPSP amplitude (mV) pre (−5 to 0 min) and post induction (25-30 min) with inhibition intact (black) or with dendritic GZ (tan). B3) Average slope (mV/ms) of the 20-80% rising phase of the EPSP pre and post induction. C1) Time course of normalized EPSP amplitude pre and post induction (grey box, t=0) in control (black) and with dendritic Gabazine (tan). C2) Time course of normalized EPSP slope, colors as in C2. C3) Normalized EPSP amplitude, and slope 30 min post induction in control (black) and with GZ (tan).

However, in neurons where stability could be verified, potentiation typically lasted for the duration of recording (45-60 min). Both raw and normalized (to baseline) EPSP amplitude and rising slope were analyzed. PNs were excluded if input resistance or membrane potential varied from baseline (±20% or ±5 mV respectively).

### Disinhibition of PN dendrites promotes LTP

With inhibition intact, the induction protocol did not induce LTP of L1b synapses. Neither EPSP amplitude (in mV, pre: 2.6 ± 0.60, post: 2.6 ± 0.51, p: 0.85, paired t-test, n=9) nor slope (in mV/ms, pre: 0.64 ± 0.20, post: 0.61 ± 0.18, p: 0.27, paired t-test, n=9) significantly differed from baseline (pre) at 30 min post induction (**Fig 1B**). Likewise, normalized EPSP amplitude and area did not differ significantly from 1 (Amplitude: 1.0 ± 0.071, p: 0.65; Slope: 0.96 ± 0.079, p: 0.65, one sample t-test, **Fig 1C, D**). However, when we focally applied the GABA_A_ receptor antagonist, Gabazine (GZ, 20 µM), to L1b (schematic, **Fig 1A**), induction significantly enhanced EPSP amplitude (mV, Pre: 1.3 ± 0.32, Post: 3.1 ± 0.95, p<0.050 WSR, n=6, **Fig 1B**) and slope (mV/ms, Pre: 0.28 ± 0.070, Post: 0.52 ± 0.12, p<0.050 WSR, n=6). Normalized EPSP amplitude and slope were significantly greater than 1 (Amplitude: 2.3 ± 0.35, p<0.05 WSR; Slope: 2.0 ± 0.27, p<0.05 WSR, **Fig 1C, D**). Consistent application of dendritic Gabazine for the duration of recording is difficult to maintain. In a some PNs, EPSP amplitudes were potentiated but unstable, these were excluded from analysis (n=5). This highlights the need for a more selective and reliable optogenetic approach to investigating disinhibition.

### Inactivation of SST-INs promotes LTP

We have previously shown that SST-interneurons strongly inhibit PN dendrites in L1B (Large, Kunz et al. 2016). Although SST-INs typically receive weak direct L1a input (Suzuki and Bekkers 2010a), SST-INs are recruited by strong TBS of L1a through the recruitment of recurrent excitation (**Fig 2A1,2)**. To investigate the influence of SST-INs on LTP induction, Archaerhodopsin (Arch) was selectively expressed in SST-INs for optical inactivation. During TBS, optical inactivation reduced SST-IN firing rate (FR) from 6.8 ± 1.2 Hz to 1.8 ± 0.4 Hz (p<0.05, WSR test, **Fig 2A2**) and enhanced depolarization in PNs (**Fig 2B**). These findings suggest that SST-interneurons mediate inhibition during TBS stimulation and could prevent associative synaptic plasticity at L1B synapses. To test this, we optically inactivated SST-INs solely during LTP induction (2 sec). EPSP amplitude and slope was significantly enhanced 30 min post induction (Amplitude (mV), Pre: 2.6 ± 0.41, Post: 4.3 ± 0.76, p: 0.0071; Slope (mV/ms), Pre: 0.55 ± 0.14, Post: 1.1 ± 0.23, p: 0.0048; n=9, paired t-test, **Fig 2C1,2, E**). Normalized EPSP amplitude and slope were significantly greater than 1 (Amplitude: 1.7 ± 0.16, p: 0.00010; Slope: 2.0 ± 0.24, p: 0.0033, one sample t-test, **Fig 2D**). EPSP traces from representative PNs pre and post induction are shown in **Fig 2E**. The normalized amplitude and slope averaged across neurons over 30 min are shown in **Fig 2F1,2**. To confirm that associative synaptic plasticity was NMDA-receptor dependent, we bath applied the NMDAR antagonist, DL-APV (10 mM) to a second set of slices. Antagonism of NMDAR prevented LTP induction despite inactivation of SST-INs. EPSP amplitude and slope 30 min post induction did not significantly differ from baseline (Amplitude (mV): Pre: 4.7 ± 0.81, Post: 4.0 ± 0.73, p: 0.21; Slope (mV/ms), Pre: 0.83 ± 0.19, Post: 0.74 ± 0.18, p: 0.26; n=10, paired t-test, **Fig 2C, black circles)**; (normalized Amplitude: 0.88 ± 0.13, p: 0.38; Slope: 0.85 ± 0.090, p: 0.28, one sample t-test. **Fig 2D**,**F black circles**). These findings suggest that inhibition by SST-INs regulates NMDA-dependent associative LTP at L1b intracortical synapses.

**Figure 2:**
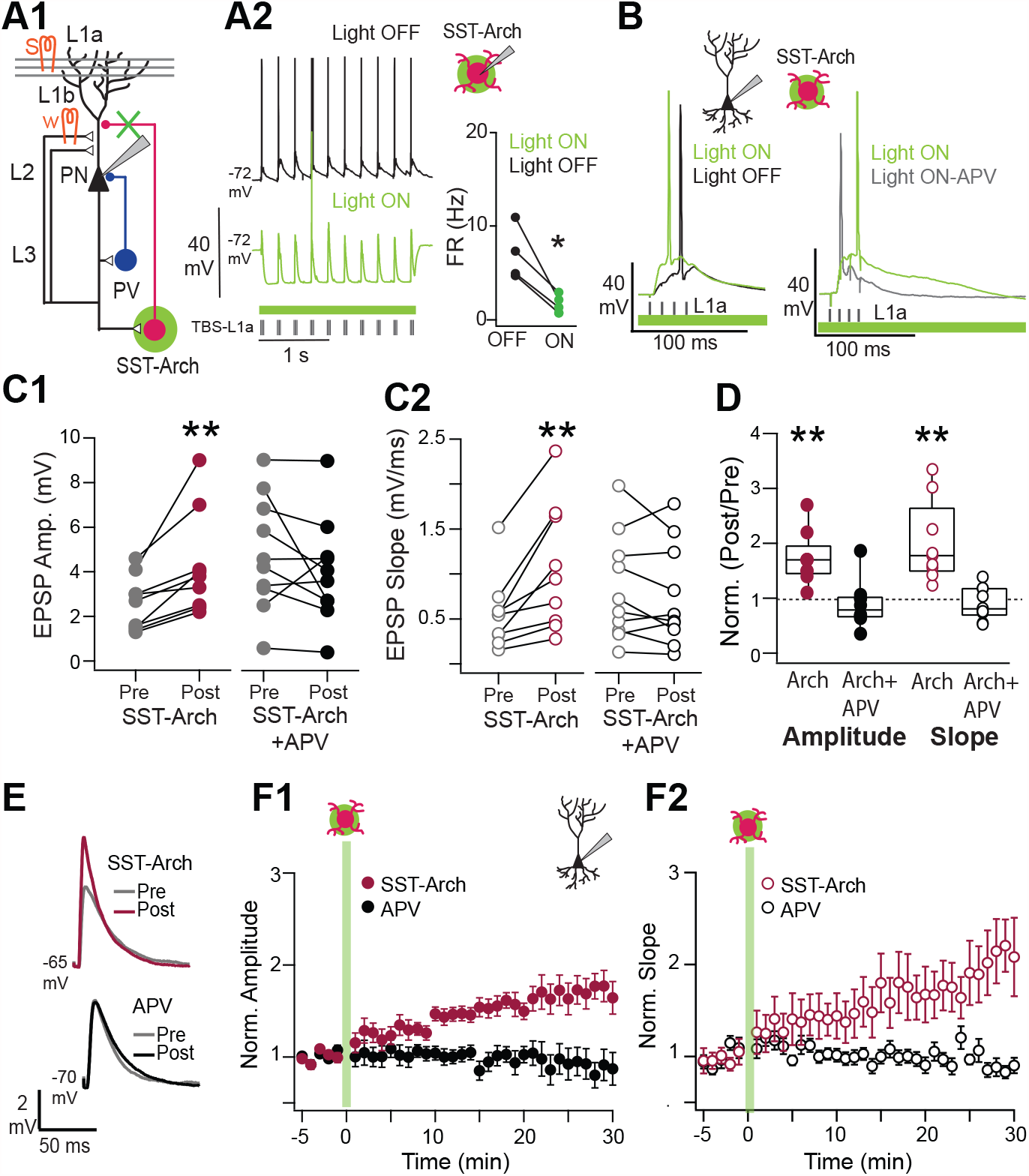
SST-interneuron inactivation promotes associative LTP. A1) Schematic: SST-INs express Archaerhodopsin (Arch). A2) SST-IN spike responses during TBS in control (black) and inactivated (green) conditions (*p<0.05, WSR test). B) PN responses for a single TBS burst. Left : Inactivation of SST-INs enhanced PN depolarization (green vs. black trace). Right: Depolarization during SST-IN inactivation (green) is reduced by APV (gray trace). C1) Average EPSP amplitude (mV) pre (−5 to 0 min) and post induction (25-30 min) with SST-inactivation (Magenta) or with SST-inactivation plus bath application of the NMDAR antagonist, APV (black). C2) Average slope (mV/ms) of the 20-80% rising phase of the EPSP pre and post induction, colors as in B2. C3) Normalized EPSP amplitude and slope following pairing with SST-IN inactivation (magenta circles). LTP is blocked by bath application APV (black). D1) Representative average traces pre (gray) and post induction in with SST-IN inactivation (magenta) or inactivation plus APV (Black). D2) Time course of normalized EPSP amplitude pre and post induction with inactivation of SST-INs (green box, t=0) in control (magenta) and with bath application of APV (black). D3) Time course of normalized EPSP slope, colors as in D2.

### Inactivation of PV-INs promotes LTP

PV-INs strongly inhibit cortical neurons. We investigated whether optogenetic inactivation of PV-INs expressing Arch also promotes associative LTP (**Fig 3A1**). PV-INs are robustly activated during TBS **(Fig 3A2)** and light inactivation significantly decreased FR (Light OFF: 7.2 ± 2.5 Hz, ON: 2.8 ± 1.7 Hz, n=5, p<0.05, WSR, **Fig 3A2, right)**. In a number of PNs, this enhanced EPSP summation **(Fig 3B1**) and/or FR consistent with somatic disinhibition. PV-IN inactivation during induction promoted LTP of EPSP amplitude and slope **(Fig 3C1,C2**, Amplitude (mV), Pre: 1.6 ± 0.34, Post: 2.4 ± 0.40, p: 0.00095; Slope (mV/ms): 0.38 ± 0.10, Post: 0.66 ± 0.18, p: 0.016; n=9, paired t-test). Normalized amplitude and slope were significantly >1 post induction (Amplitude: 1.6 ± 0.16, p: 0.0063; Slope: 1.8 ± 0.15, p: 0.00061, one sample t-test **Fig 3C3, D**). Antagonism of NMDAR prevented LTP induction mediated by inactivation of PV-INs. EPSP amplitude and slope post induction did not significantly differ from baseline (Amplitude (mV): Pre: 3.3 ± 0.45, Post: 3.0 ± 0.37, p: 0.48; Slope (mV/ms), Pre: 0.44 ± 0.082, Post: 0.52 ± 0.10, p: 0.74; n=8, paired t-test, **Fig 3C, black circles)**; (normalized Amplitude: 0.92 ± 0.076, p: 0.65; Slope: 1.0 ± 0.13, p: 0.56, one sample t-test. **Fig 3C,D black circles**). Thus, PV-INs can also regulate the induction of NMDA-dependent associative LTP at L1b intracortical synapses.

**Figure 3:**
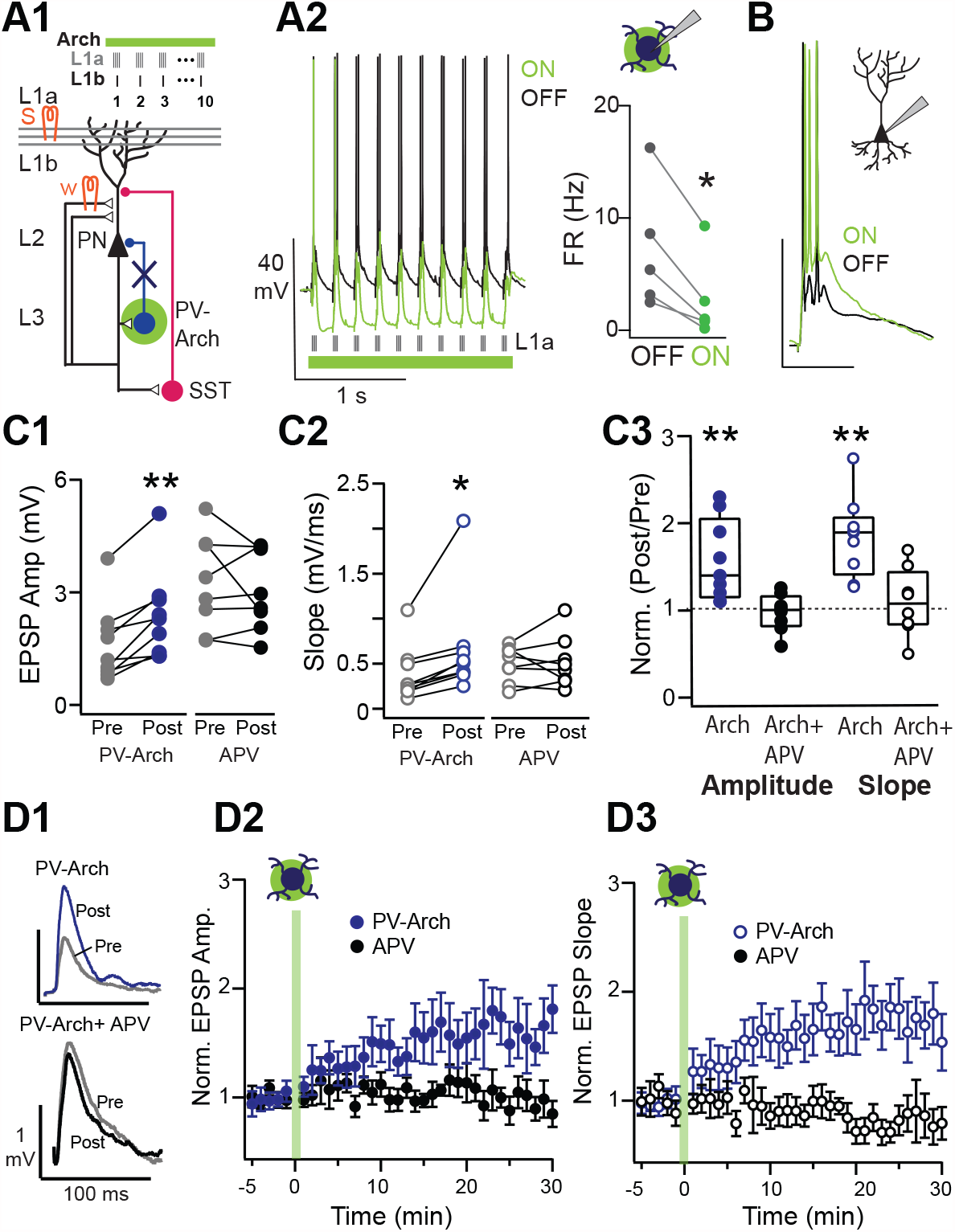
PV-interneuron inactivation during induction promotes associative LTP. A1) Schematic: PV-INs express Archaerhodopsin (Arch). A2) PV-IN spike responses during TBS in control (black) and inactivated (green) conditions. B) Representative PN responses for a single TBS for control (black) versus PV-IN inactivation (green). C1) Average EPSP amplitude (mV) pre (−5 to 0 min) and post induction (25-30 min) with PV-inactivation (Blue) or with PV-inactivation plus bath application of the NMDAR antagonist, APV (black). C2) Average slope (mV/ms) of the 20-80% rising phase of the EPSP pre and post induction, colors as in B2. C3) Normalized EPSP amplitude and slope following pairing with PV-IN inactivation (blue circles) or with inactivation plus bath application APV (black). D1) Representative average traces pre (gray) and post induction in with PV-IN inactivation (blue) or inactivation plus APV (Black). D2) Time course of normalized EPSP amplitude pre and post induction with inactivation of PV-INs (green box, t=0) in control (blue) and with bath application of APV (black). D3) Time course of normalized EPSP slope, colors as in D2.

### Inhibition of interneurons by VIP-INs

If both SST-IN and PV-INs can inhibit LTP induction, the question remains as to which circuits might provide a means for disinhibition and the promotion of LTP. In other cortices, VIP interneurons inhibit SST-INs and PV-INs (Pfeffer, Xue et al. 2013, Pi, Hangya et al. 2013). Although there are numerous VIP-INs in piriform cortex (Suzuki and Bekkers 2010b), their targets have not been identified. We crossed VIP-cre mice with Ai32 mice to express channelrhodopsin (ChR2) in VIP-INs. We then used light activation to drive single action potentials in VIP-INs while recording potential postsynaptic targets. Typically, VIP-cre mice are crossed with mice that express GFP in SST-INs (GIN-mice) or PV-INs (G42-mice). However, these lines sparsely label SST and PV-INs in APC (Large, Kunz et al. 2016). Instead, we used intrinsic properties to identify putative (p) SST-INs and pPV-INs according criteria determined from recordings in genetically identified SST-INs and PV-INs in APC (Large, Kunz et al. 2016)(see methods). Interneurons that could not be confidently identified were excluded from analysis.

VIP-INs were activated by brief blue light pulses (5 ms) and IPSCs were recorded in voltage clamp with either Cs-Gluconate-Qx internal (IPSCs, +30 mV) or K-Gluconate-Qx internal (IPSCs, 0 mV). IPSC recordings were most reliable with Cs-Glu-Qx and the data presented (**Fig 4)** are from this condition. VIP-INs inhibited nearly all recorded pSST-INs (86%), most pPV-INs (90%) and PNs (88%). Although high connectivity with SST and PV cells was expected, this was unexpected for PNs (Pfeffer, Xue et al. 2013). VIP-INs strongly inhibited pSST-INs (IPSC amplitude: 303 ± 57 pA, n=19). However, despite a high probability of connection, VIP-INs very weakly inhibited pPV-INs (35 ± 5.8 pA, n=9, p: 0.0008) and PNs (61 ± 14 pA, p: 0.001 n=14, ANOVA, **Fig 4A1**). Qualitatively similar results were found with K-Glu-Qx, however IPSC amplitude and connection probability were considerably lower (% connected, amplitude, # connected: pSST: 73%, 47 ± 14 pA, n=19; pPV: 73%, 31 ± 7.6 pA, n=11; PN: 29%, 21 ± 7.0 pA, n=4).

**Figure 3:**
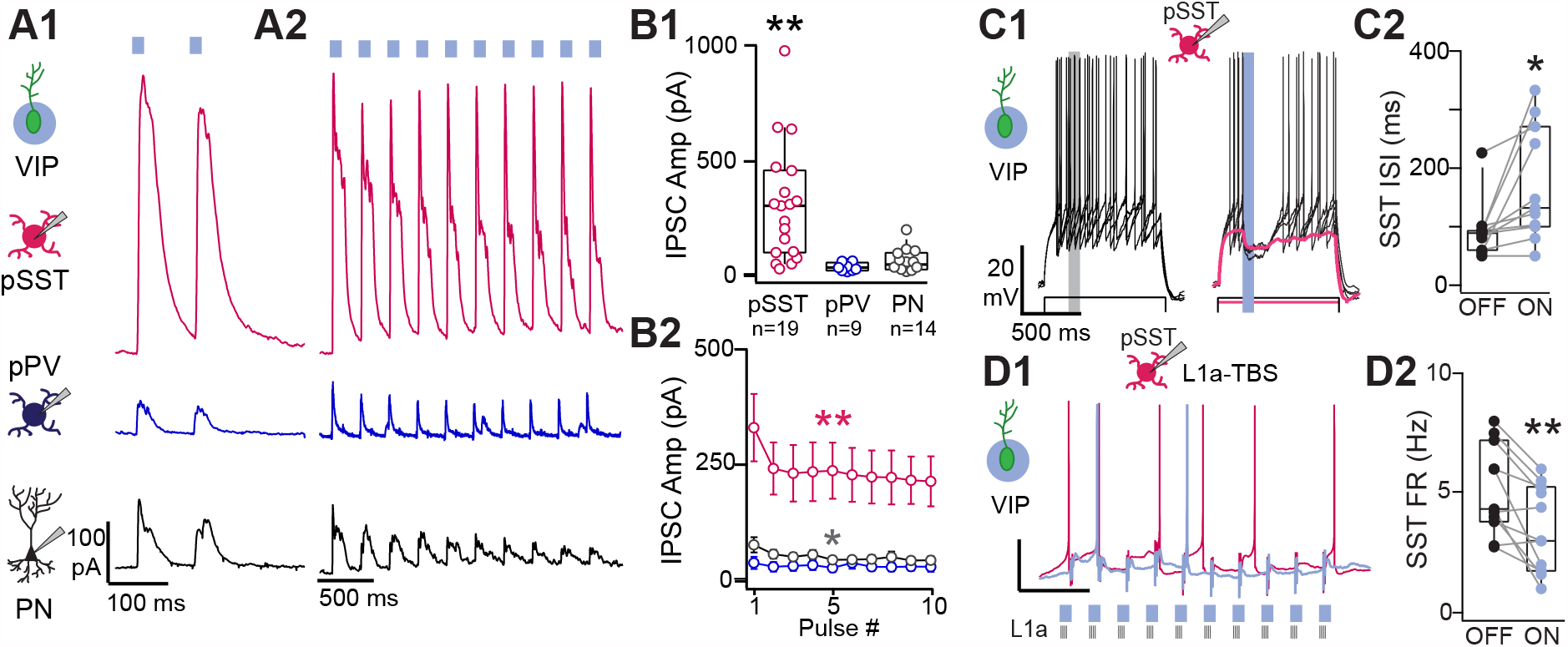
Inhibition by VIP-interneurons in piriform cortex. A1) VIP-INs express ChR2. Optically evoked IPSCs were recorded in putative(p) pSST-INs (magenta), pPV-INs (blue) and PNs (black). A2) Responses of the same neurons (A1) to ten light pulses (100 ms duration, 4 Hz). B1) IPSC amplitude was stronger in pSST-INs versus pPV-INs (*p:0.014) or PNs (*p:0.042, ANOVA). B2) In pSST-INs, IPSC amplitude diminishes by the 5th pulse of theta stimulation (**p: 0.009, paired t-test). C1) IPSCs from VIP-INs delay pSST-IN spike responses during suprathreshold depolarization (4 overlaid traces) Left: Control, Right: activation of VIP-INs (100 ms pulse, blue). Magenta trace: VIP-IN mediated IPSC during subthreshold depolarization. C2) Interspike interval (ISI) was significantly increased during optical activation of VIP-INs (blue circles, p: 0.016, paired t-test, n=11) compared to light off trials (black circles). D1) Spike responses in pSST-INs during TBS in control (magenta trace) and during pulsed light (blue trace). D2) pSST-IN firing rates decreased during TBS on light trials (blue) versus control (black circles, p:0.002, paired t-test, n=11).

We find that VIP-INs typically show strongly adapting spike responses within the first 500 ms of suprathreshold depolarizations by current steps. For this reason, long light pulses (2 s) during LTP induction would likely result in significant adaptation of VIP-INs spike responses. Instead, we drove VIP-INs with shorter light pulses (100 ms) delivered at theta frequency (10 pulses, **Fig 4A2**). This pulsed-light stimulation evoked strong and relatively sustained IPSCs in pSST-INs (**Fig 4B2 top**). Overall, IPSC strength decreased by only ∼30% by the 5^th^ pulse in all cell types then stabilized (SST: 31 ± 10%, p: 0.002; PV: 31± 5%, p: 0.06; PC: 28 ± 12%, p: 0.02, paired t-test, **Fig 4B2**).

To investigate how VIP-IN mediated inhibition could influence action potential activity we depolarized target neurons to near or suprathreshold membrane potentials in current clamp. In pSST-INs activation of VIP-INs evoked strong IPSPs (2.02 ± 0.38 mV, n=18, **Fig 4C1**). These IPSPs were sufficient to delete or delay spikes by an average of 77 ± 26 ms (p: 0.016, n=11, paired t-test) compared to non-light trials (**Fig 4C1,3 left**). Finally, pSST-IN spike responses evoked by TBS were significantly decreased by pulsed activation of VIP-INs compared to non-light trials (OFF: 5.0 ± 0.6 Hz, ON: 3.5 ± 0.5, p: 0.002, n=11, paired t-test, **Fig 4C**). We could not record IPSPs in PNs (n=7) or pPV-INs (n=4) likely due to a combination of low input resistance and weak VIP-mediated IPSCs. Taken together, these findings suggest that VIP-INs could strongly influence pSST-INs during LTP induction but would have considerably less impact on pPV-IN or PNs.

### Activation of VIP-INs promotes LTP

VIP-INs are numerous in APC and we find that VIP-INs inhibit pSST-INs. VIP-to-SST inhibition is a candidate circuit motif for dendritic disinhibition of PNs that could promote LTP. So, why isn’t LTP induced without antagonism of inhibition? We recorded VIP-INs during TBS stimulation of L1a. We found that in contrast to SST-INs or PV-INs, VIP-INs were only weakly driven to fire action potentials by TBS (0.7 ± 0.6 Hz, n=6, Fig 5A2). This lack of VIP-IN recruitment is consistent with the inability to induce LTP under control conditions (**Fig 1B**). However, light activation of ChR2+ VIP-INs enhanced FR during TBS (14 ± 4.0 Hz, p<0.05, WSR test **Fig 5A2**). Activating VIP-INs also enhanced EPSP summation in PNs during TBS **(Fig 5B)**. Activation of VIP-INs during induction lead to robust LTP of EPSP amplitude and slope (**Fig 5C1,C2**, amplitude (mV), Pre: 2.1 ± 0.50, Post: 3.4 ± 0.48, p<0.02; slope (mV/ms) Pre: 0.47 ± 0.081, Post: 0.76 ± 0.072, p<0.02, n=7, WSR). Normalized amplitude and slope were significantly >1 post induction (Amplitude: 2.0 ± 0.33, p<0.02; Slope: 1.9 ± 0.32, p<0.02, WSR, **Fig 5C3**). Antagonism of NMDARs prevented LTP induction mediated by activation of VIP-INs. EPSP amplitude and slope post induction did not significantly differ from baseline (Amplitude (mV): Pre: 6.0 ± 1.6, Post: 5.5 ± 1.5, p>0.05; Slope (mV/ms), Pre: 1.2 ± 0.23, Post: 1.1 ± 0.28, p: >0.05; n=5, WSR, **Fig 5C,D black circles)**; (normalized Amplitude: 0.93 ± 0.072, p>0.05; Slope: 0.99 ± 0.17, p>0.05, WSR. **Fig 5C,D black circles**). Altogether, these findings support a VIP-SST-PN disinhibitory circuit that gates associative LTP at intracortical synapses.

**Figure 5:**
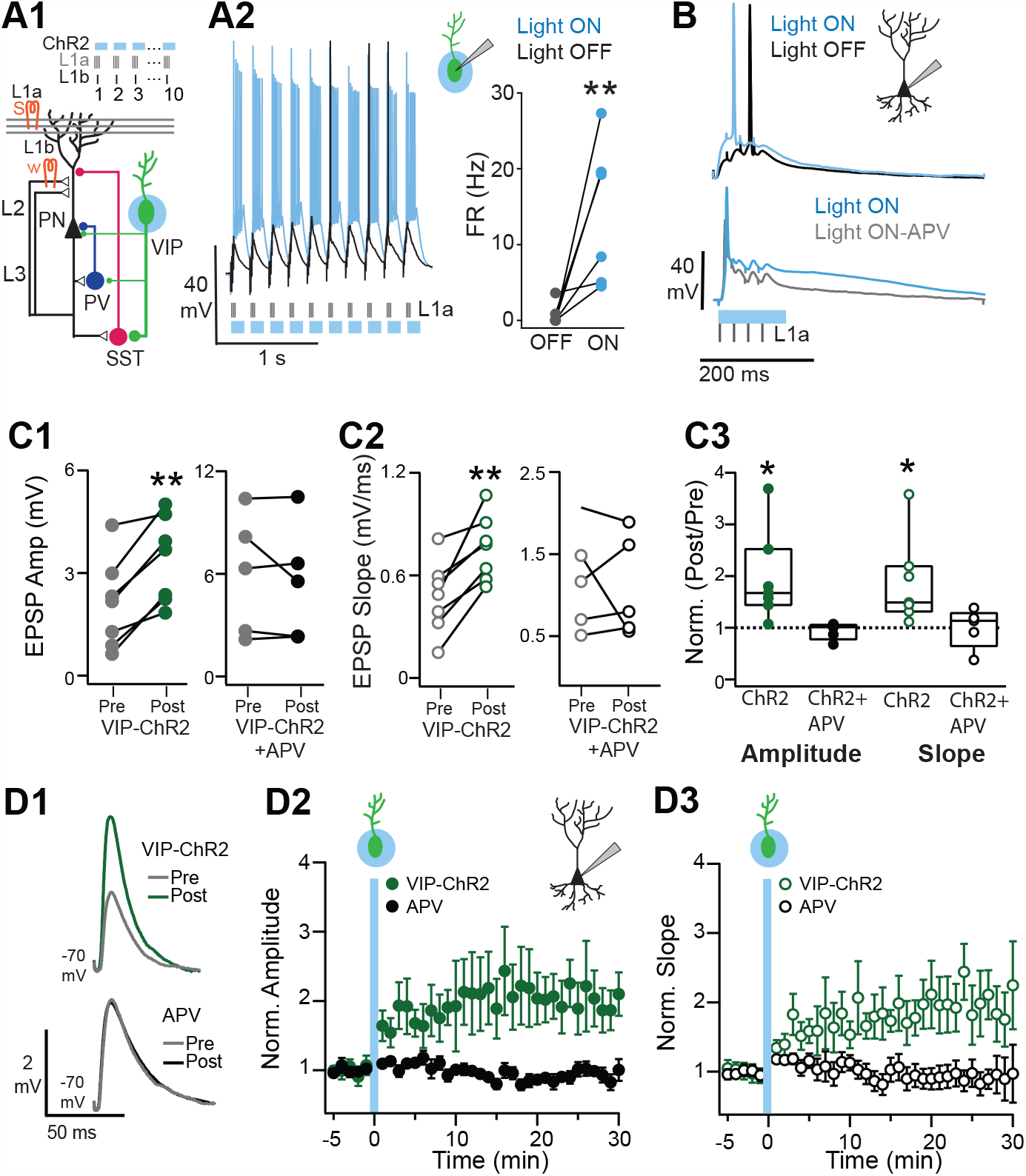
VIP-interneuron activation promotes associative LTP. A1) Circuit schematic: VIP-INs express ChR2 and were activated using theta pulsed light during L1a+L1b pairing. A2) VIP-IN responses during TBS without (black) and with light (blue). FRs increase during pairing with light (blue circles, p<0.05, WSR, n=7). B) Top: Activation of VIP-INs enhanced PN depolarization during TBS stimulation (blue vs. black trace). Bottom: PN depolarization during VIP-IN activation (blue trace) is reduced by APV (gray trace). C1) Average EPSP amplitude (mV) pre (−5 to 0 min) and post induction (25-30 min) with VIP-IN activation (green) or with VIP-IN activation plus bath application of APV (black). C2) Average slope (mV/ms) of the rising phase of the EPSP pre and post induction, colors as in C1. C3) Normalized EPSP amplitude and slope following pairing with VIP-IN activation (green circles) or with activation plus APV (black). D1) Representative average traces pre (gray) and post induction in with VIP-IN activation (green) or activation plus APV (Black). D2) Time course of normalized EPSP amplitude pre and post induction with activation of VIP-INs (blue box, t=0) in control (green) and with APV (black). D3) Time course of normalized EPSP slope, colors as in D2.

## Discussion

It has been hypothesized that odor ensembles are formed through associative plasticity of recurrent excitatory synapses between co-activated pyramidal neurons (Haberly 2001, Wilson and Sullivan 2011). These intracortical synapses are stronger in animals that have learned an olfactory discrimination task (Saar, Grossman et al. 2002, Saar, Reuveni et al. 2012) and the induction of associative LTP *in vitro* is occluded in animals that have learned tasks (Lebel, Grossman et al. 2001). This capacity for enhancement as well as highly recurrent excitation (Poo and Isaacson 2011) necessitates strong inhibition to regulate neural activity (Luna and Schoppa 2008, Poo and Isaacson 2011, Bolding and Franks 2018) and synaptic plasticity (Kanter and Haberly 1993, Kanter, Kapur et al. 1996). Our present findings are consistent with a VIP->SST->PN circuit that transiently disinhibits PN dendrites to promote synaptic plasticity during odor learning while overall inhibition remains intact.

Associative LTP at intracortical synapses within APC is well-characterized (Stripling, Patneau et al. 1988, Kanter and Haberly 1993, Poo and Isaacson 2007). TBS provides strong afferent excitation that depolarizes apical PN dendrites and promotes NMDAR dependent potentiation of co-activated intracortical synapses. However, strong TBS also recruits strong inhibition and LTP is rarely induced in the absence of GABA_A_R antagonists (Kanter and Haberly 1993). We have reproduced previous findings that LTP is gated by dendritic application of Gabazine (Kanter, Kapur et al. 1996, Kumar, Schiff et al. 2018). Further, we have shown that dendritic application of Gabazine blocks inhibition mediated by SST-INs (Large et al., 2016). In the present study, we demonstrate that inactivation of SST-INs at an opportune moment during TBS, is sufficient to promote LTP. Thus, a circuit mechanism that inhibits SST-INs during LTP induction could gate synaptic plasticity. Throughout the cortex, VIP-INs inhibit SST-INs and potentially disinhibit PNs (Lee, Kruglikov et al. 2013, Pfeffer, Xue et al. 2013, Karnani, Jackson et al. 2016). We show that VIP-INs strongly inhibit pSST-INs and weakly inhibit pPV-INs as well as PNs. Moreover, optogenetic activation of VIP-INs decreases pSST-IN spike responses during TBS. Finally, activation of VIP-INs specifically during TBS is sufficient gate associative LTP. Based on these findings, we propose that a VIP->SST->PN disinhibitory circuit gates intracortical synaptic plasticity in APC.

However, it is relevant to discuss three additional circuits that are potentially recruited with VIP-IN activation. For example, VIP-INs could also gate plasticity through a VIP-PV-PN disinhibitory circuit. Although optogenetic inactivation of PV-INs promotes LTP, we predict this circuit motif contributes minimally to associative LTP induction. VIP-INs very weakly inhibit pPV-INs, which is likely insufficient to prevent PV-IN spiking during TBS. Conversely, SST-INs strongly inhibit PV-INs (Pfeffer, Xue et al. 2013, Xu, Jeong et al. 2013, Large, Kunz et al. 2016) and inactivation of SST-INs in APC increases pPV-IN activity (Sturgill and Isaacson, 2015). Instead of disinhibiting the PN, a VIP-SST-PV circuit could transiently increase somatic PV-IN inhibition to compensate for a loss of SST mediated inhibition at the dendrite and/or soma. Finally, VIP-INs also inhibit PNs directly, possibly through basket like synapses on the soma (Suzuki and Bekkers 2010a). As with VIP-PV synapses, this VIP-PN inhibitory circuit is also weak and its role is unknown. Importantly, recruitment of these additional circuits does not impair LTP induction with VIP-IN activation. Rather, the interplay of these circuits suggests a shift in inhibition from the dendrite to soma that may stabilize neural responses while opening a window for plasticity.

Our study complements a recent study in somatosensory cortex that used chemogenetics to inhibit VIP-INs or SST-INs and prevent or promote LTP induction respectively (Williams and Holtmaat 2019). Together, these two studies support a common VIP->SST->PN disinhibitory circuit motif regulates synaptic plasticity across cortical areas. A benefit of the present study is that by using optogenetics, we were ability to isolate the influence of disinhibition to a brief time window (∼2 s) when afferent and intracortical pathways are co-activated. This suggests that transient dendritic disinhibition is sufficient to gate the cascade of intracellular mechanisms underlying synaptic enhancement. Thus, circuit mechanisms that provide a means for VIP cells to be activated in conjunction with odor sampling could play a key role in odor learning and ensemble formation.

Interestingly, VIP-INs are not recruited by pairing afferent and intracortical stimulation, this may underlie the inability induce associative LTP with inhibition intact (**Fig 1**). At present, it is not known how VIP-INs in APC are recruited. In other cortices, VIP-IN activity is enhanced by arousal, locomotion, task engagement or reward (Lee, Kruglikov et al. 2013, Pi, Hangya et al. 2013, Fu, Tucciarone et al. 2014, Jackson, Ayzenshtat et al. 2016), either through additional excitatory drive from other cortical areas (Lee, Kruglikov et al. 2013, Williams and Holtmaat 2019) or through neuromodulation (Porter, Cauli et al. 1999, Lee, Hjerling-Leffler et al. 2010, Alitto and Dan 2012, Kuchibhotla, Gill et al. 2017, Pronneke, Witte et al. 2019). Direct excitatory inputs to VIP-INs would have to be well-timed with respect to sensory stimulation. Conversely, neuromodulation could raise the excitability of VIP-INs such that afferent and intracortical drive are sufficient to drive VIP-INs with appropriate timing. The APC receives input from higher cortices including orbitofrontal cortex (Illig 2005) as well as neuromodulatory centers (Zaborszky, Carlsen et al. 1986, Linster, Wyble et al. 1999). Both of these pathways have been implicated in olfactory learning and plasticity (Patil, Linster et al. 1998, Patil and Hasselmo 1999, Linster, Maloney et al. 2003, Li, Luxenberg et al. 2006, Chapuis and Wilson 2013, Cohen, Wilson et al. 2015, Strauch and Manahan-Vaughan 2018). Future studies are needed to ascertain the potential links between descending excitation and/or neuromodulation, the recruitment of VIP-INs, and the gating of synaptic plasticity during olfactory processing.

Though many previous studies have focused on the influence of disinhibitory circuits on FR or behavior, our study highlights a role for disinhibition in the dynamics that gate long-term plasticity. Specifically, our findings suggest that the VIP-SST-PN circuit motif plays a central role in gating synaptic plasticity in PN dendrites. Additional circuit mechanisms mediated through VIP-INs that regulate PV-INs or PNs directly could serve a stabilizing role. These findings demonstrate the challenge of the delineating roles for the individual circuit motifs that are nested in complex neural networks. It remains to be determined if these mechanisms work in concert during afferent and recurrent olfactory processing.

## Notes

### Competing Interest Statement

The authors have declared no competing interest.

### Summary of Updates

We have updated data, analysis and all figures in accordance with suggestions from other scientists.

